# Tissue-dependent nature of plant susceptibility: a comparative pathogenicity study of the systemic phytopathogen *Ralstonia pseudosolanacearum* in eggplant and tomato seedlings through root and leaf

**DOI:** 10.1101/2024.11.25.625158

**Authors:** Shuvam Bhuyan, Monika Jain, Lakhyajit Boruah, Tana Sun Tara, Shuhada Begum, Lukapriya Dutta, Shubhra Jyoti Giri, Tarinee Phukan, Kristi Kabyashree, Manabendra Mandal, Suvendra Kumar Ray

**Affiliations:** Department of Molecular Biology & Biotechnology, Tezpur University, Tezpur–784028, Assam, India; Plant-Microbe Interactions Laboratory, National Institute of Plant Genome Research, Aruna Asaf Ali Marg, New Delhi–110067, India

**Keywords:** *Ralstonia pseudosolanacearum*, virulence, pathogenicity, plant response, tomato, eggplant

## Abstract

*Ralstonia pseudosolanacearum* causes a lethal bacterial wilt disease in many plant species, posing significant economic challenges. Although tomato has been a primary model host for investigating the pathogenicity and systemic infection of this bacterium, this manuscript presents a comparative pathogenicity study between two closely related solanaceous hosts, tomato and eggplant, revealing differential host responses to the same pathogen, specifically through leaves and roots. Interestingly, eggplant seedlings exhibited a significantly higher susceptibility to cotyledon leaf inoculation than tomato seedlings. In the case of tomato, a few seedlings escaped wilting (called escapees), which was usual at a high pathogen load, unlike the case of eggplant seedlings where escapees could not be observed even at a 100-fold lower pathogen concentration. The greater susceptibility of eggplant was further demonstrated by performing both single- and double-leaf inoculations in the seedlings. Surprisingly, root inoculations resulted in a significantly lower mortality for eggplant than for tomato seedlings. The contrasting susceptibility between the two hosts regarding root and leaf regions indicates the tissue-dependent nature of susceptibility. The study underscores the value of employing multiple host species to uncover new insights into pathogen behaviour and host-pathogen interactions.

## 1. Introduction

Bacterial wilt caused by the Gram-negative bacterial species complex of *Ralstonia solanacearum, R. pseudosolanacearum* and *R. syzygii* was reported more than a century ago [1,2]. The pathogen exhibits a broad host range, capable of infecting over 200 plant species belonging to 53 botanical families [3]. Unlike many plant-pathogenic bacteria that are tissue-specific, *R. pseudosolanacearum* systemically infects the entire host plant, eventually leading to wilting [4–8]. Under laboratory conditions, tomatoes have been used as the major model host to study their pathogenicity. Therefore, reports are scanty regarding its comparative pathogenicity among multiple hosts, even though it is a broad host range pathogen. Despite extensive global research, there remain several fundamental questions regarding its systemic infection that require more comprehensive investigation. Some of these questions include: (i) How does the pathogen cross the root-shoot junction while moving from root to shoot during infection? (ii) Is systemic colonization a prerequisite for causing wilting? If so, then how does the pathogen transition from the colonization phase to the pathogenicity phase? (iii) Why is the disease phenotype observed in the shoot region but not in the root, even though the pathogen naturally enters the plant through the root? (iv) Why do a few of the inoculated tomato plants not exhibit disease symptoms (known as "escapees") even in the presence of the pathogen inside these plants? (v) What enables the pathogen to adapt to diverse hosts? The underlying mechanisms behind this behavior remain unclear.

In general, *R. pseudosolanacearum* pathogenicity is studied using soil drenching and stem-inoculation methods in 1-2 months old grown-up tomato plants to mimic natural infection. However, in our recent approach, 6- to 7-day-old early cotyledon stage seedlings of tomato and eggplant have been demonstrated as model hosts to study *R. pseudosolanacearum* pathogenicity [6–8]. Seedlings being small in size, this change allows us to conduct large-scale pathogenicity studies under controlled laboratory conditions. Our recent studies in tomato and eggplant seedlings have revealed unique pathological phenotypes that had not been previously reported in seedling inoculation experiments. Notably, the symptom of stem softening and translucency was observed more frequently in root-inoculated eggplant seedlings, highlighting a distinct difference in the bacterium’s virulence between tomato and eggplant seedlings [9]. In addition, using sensitive as well as resistant tomato cultivars in their early growth stage, it has been demonstrated that the pathogen movement usually stops in the resistant host [10].

Considering the susceptibility of both tomato and eggplant seedlings towards *R. pseudosolanacearum* infection, we carried out a comparative analysis of its pathogenicity between these two phylogenetically close solanaceous crops: tomato (*Solanum lycopersicum*) and eggplant (*Solanum melongena*). Being dicotyledonous, each seedling of tomato and eggplant possesses a pair of cotyledon leaves that provided us with the avenue to study and compare the pathogenicity of *R. pseudosolanacearum* in seedlings inoculated at single site (single cotyledon leaf) vs double site (both the cotyledon leaves). By employing single- and double-leaf inoculation modes, we could distinctly demonstrate the higher susceptibility of eggplant in comparison to tomato seedlings. It is interesting to observe that though eggplant seedlings are more susceptible than tomato seedlings to *R. pseudosolanacearum* infection by leaf inoculation, the reverse is true in the case of root inoculation. The observation of the differential nature of susceptibility between the root and the leaf could only be possible because of the comparative pathogenicity study between the two hosts. Therefore, the magnitude of susceptibility of a plant to a systemic pathogen may vary depending on the tissue involved. Furthermore, this research also underscores the advantages of using two different host plants to study pathogenicity.

## 2. Methods

### (a) Bacterial strain and growth conditions

The cells of *R. pseudosolanacearum* F1C1 were cultivated in BG broth, which included 1 % peptone (Hi-Media), 0.1 % yeast extract (Hi-Media, Mumbai, India), and 0.1 % casamino acids (SRL, New Mumbai, India), along with the supplementation of 0.5 % glucose (Hi-Media) at 28 °C. In the case of solid medium, 1.5 % agar (Hi-Media) was incorporated into the BG broth [11].

### (b) Germination of seeds and growth of seedlings

We employed seeds of the tomato variety Pusa Ruby (Durga Seeds, U.P., India) and the eggplant variety Dev-Kiran (614) (Tokita, Karnataka, India) for this investigation. The seeds were germinated using the protocols of Kumar *et al.* [6]; Singh *et al.* [7]; and Phukan *et al.* [8] with the following modifications. Seeds of tomato and eggplant were subjected to a soaking process: the tomato seeds were immersed in sterile distilled H_2_O for 24 h at room temperature, and the eggplant seeds were immersed for 48 h at 4 °C. Subsequently, the seeds were to be sown to initiate germination. In all our earlier experiments involving seedlings, each batch of seeds was allowed to germinate on sterile tissue paper and an absorbent cotton-based seedbed. Our observations while using this seedbed setup for germinating seedlings were such that while the radicle from the germinated tomato seeds could penetrate the tissue paper of the seedbed and grow, the eggplant radicles failed to do so, eventually leading its roots to dry out. This suggests a strong ability of penetration to the tissue paper by the tomato root, which is lacked by the root of the eggplant seedlings. Therefore, for our comparative study, we used sterile soil as the seedbed for sowing seeds and germination.

To retain moisture, the containers with sown seeds were covered with plastic bags and kept in the dark at ambient temperature for 72 h. Following this period, the plastic covers were removed, and the sprouted seeds were transferred to a growth chamber (Orbitek, Scigenics, India) maintained at 28 °C, 12 h photoperiod and a relative humidity of 80 %. Watering of the sprouted seeds was carried out at regular intervals of 12 h to compensate for the loss of water due to transpiration and evaporation. In accordance with different experiments, the 7-day-old cotyledon stage seedlings were either transferred to 1.5 mL microfuge tubes containing sterile dH_2_O or 70 mL paper cups containing 50 g autoclaved soil for performing the infection assays. However, when transferring the seedlings to 1.5 mL microfuge tubes, the roots were first washed with sterile dH_2_O to remove the adhering soil.

### **(c)** Preparation of bacterial inoculum for virulence assays

The freshly cultured *R. pseudosolanacearum* F1C1 was introduced into 10 mL of BG broth and placed in a shaking incubator (Orbitek, Scigenics, India) maintained at 28 °C, 150 rpm for 24 h. Following this incubation, the bacterial suspension was subjected to centrifugation at 4000 rpm at 4 °C for 15 min (5804R, Eppendorf, Germany). The resulting pellet was rinsed with sterile distilled H_2_O and then reconstituted in an equivalent volume of sterile distilled H_2_O. Notably, the concentration of the bacterial suspension was adjusted to ∼ 10^9^ CFU/mL (OD_600_ is 1.0). For preparing subsequent 10-fold dilutions of the 10^9^ CFU/mL bacterial suspension, the serial dilution method was employed.

### (d) Virulence assays in tomato and eggplant seedlings

Virulence assays were conducted using 7-day-old cotyledon stage tomato and eggplant seedlings through the leaf inoculation method, as described in the works of Kumar et al. (2017) and Phukan et al. (2019). For leaf inoculation, the seedlings were removed from the soil bed and their roots were cleansed with sterile distilled H_2_O, and then placed into 1.5 mL microfuge tubes filled with sterile dH_2_O. Following this, in accordance with different experiments, approximately one-third of the cotyledon leaves were carefully excised using sterile scissors. The scissors were dipped in the bacterial inoculum for inoculation, and depending on the experimental setup, either one or both leaves were inoculated with the bacterium. In experiments where only one leaf was inoculated, the other leaf was also wounded with sterile scissors, although it was not exposed to the bacterial inoculum (Fig. 1).

**Figure 1.**
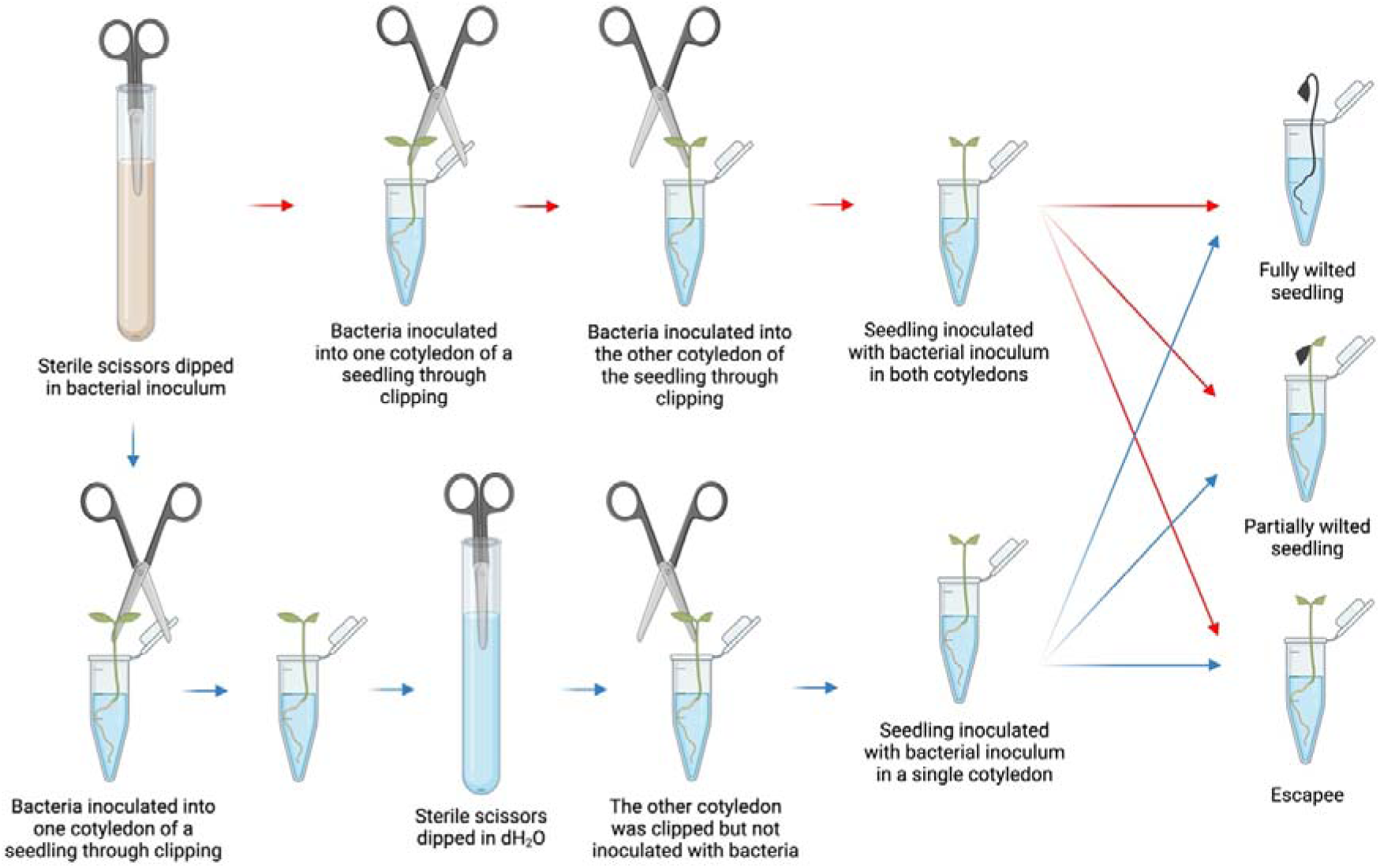
Schematic diagram of single- and double-leaf inoculation in seedlings. Created with Biorender.com.

For root inoculation, the roots of the seedlings were first dipped in the bacterial inoculum, then the seedlings were transferred to 1.5 mL microfuge tubes. After an interval of 2-5 mins sterile dH_2_O was added to it. It’s important to note that the roots of the seedlings were rinsed with sterile distilled H_2_O to remove adhering soil before being inoculated. In all experimental setups, each study group comprised 40 seedlings and the mortality of these seedlings was recorded from the day of inoculation onwards till the 10 days post inoculation (DPI) at every 24 h interval.

### **(e)** Statistical analysis

The two-sample F-tests and two-sample two-tailed T-tests for an equal variance were performed on MS-Excel to analyze the growth parameters of the seedlings. The Kaplan-Meier survivability curve was generated, and log-rank tests were conducted in R to assess the virulence of *R. pseudosolanacearum* F1C1 within its host seedlings. The R script used for this analysis has been presented in SFig. 1.

## 3. Results

### (a) Eggplant seedlings exhibit higher susceptibility to *R. pseudosolanacearum* pathogenicity when inoculated through the leaves than tomato seedlings

We studied the virulence of *R. pseudosolanacearum* F1C1 in the cotyledon stage seedlings of tomato as well as eggplant at different pathogen concentrations such as 10^9^ CFU/mL, 10^8^ CFU/mL and 10^7^ CFU/mL by the double leaf-clip inoculation. Scoring of the disease progression and seedling mortality from 0 to 10 days post-inoculation (DPI) was done (Fig. 2 & 3). In the case of eggplant, 100 % seedling mortality was observed in all three pathogen concentrations by 10 DPI. Mortality of the seedlings was initiated on 3 DPI and reached its maximum on 6 DPI. Comparative statistical analysis through log-rank tests revealed no significant difference (p-value ≥ 0.05) in the intensity of virulence across the three different pathogen concentrations of 10^9^ CFU/mL, 10^8^ CFU/mL and 10^7^ CFU/mL. In the case of tomato, seedling mortality was observed on 3 DPI, and the disease progression continued till 9 DPI. But in all the concentrations a few seedlings were found to escape the disease, which were called ‘escapees’. The percentage of escapees was 37.5 %, 25 % and 22.5 %, respectively concerning pathogen concentrations of 10^7^ CFU/mL, 10^8^ CFU/mL and 10^9^ CFU/mL. The mortality of seedlings inoculated at three different pathogen concentrations was not the same, but the disease progression was similar (p-value ≥ 0.05), i.e., no significant difference in the intensity of virulence across the three different pathogen concentrations was observed.

**Figure 2.**
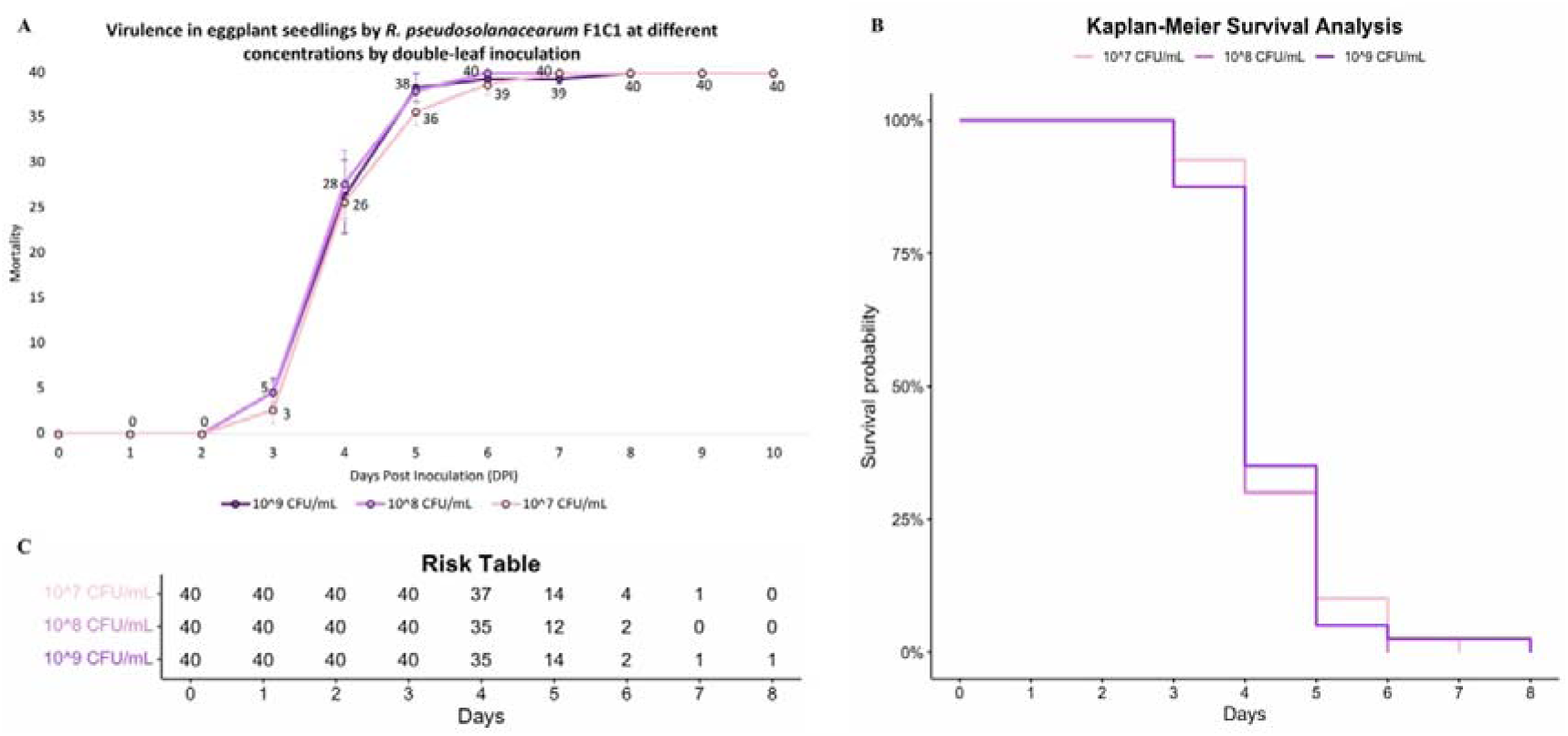
Double-leaf inoculation of higher concentrations (10^9^, 10^8^ and 10^7^ CFU/mL) of *R. pseudosolanacearum* F1C1 in eggplant seedlings. **A.** Line-graph. **B.** Kaplan-Meier survival analysis. **C.** Risk Table.

**Figure 3.**
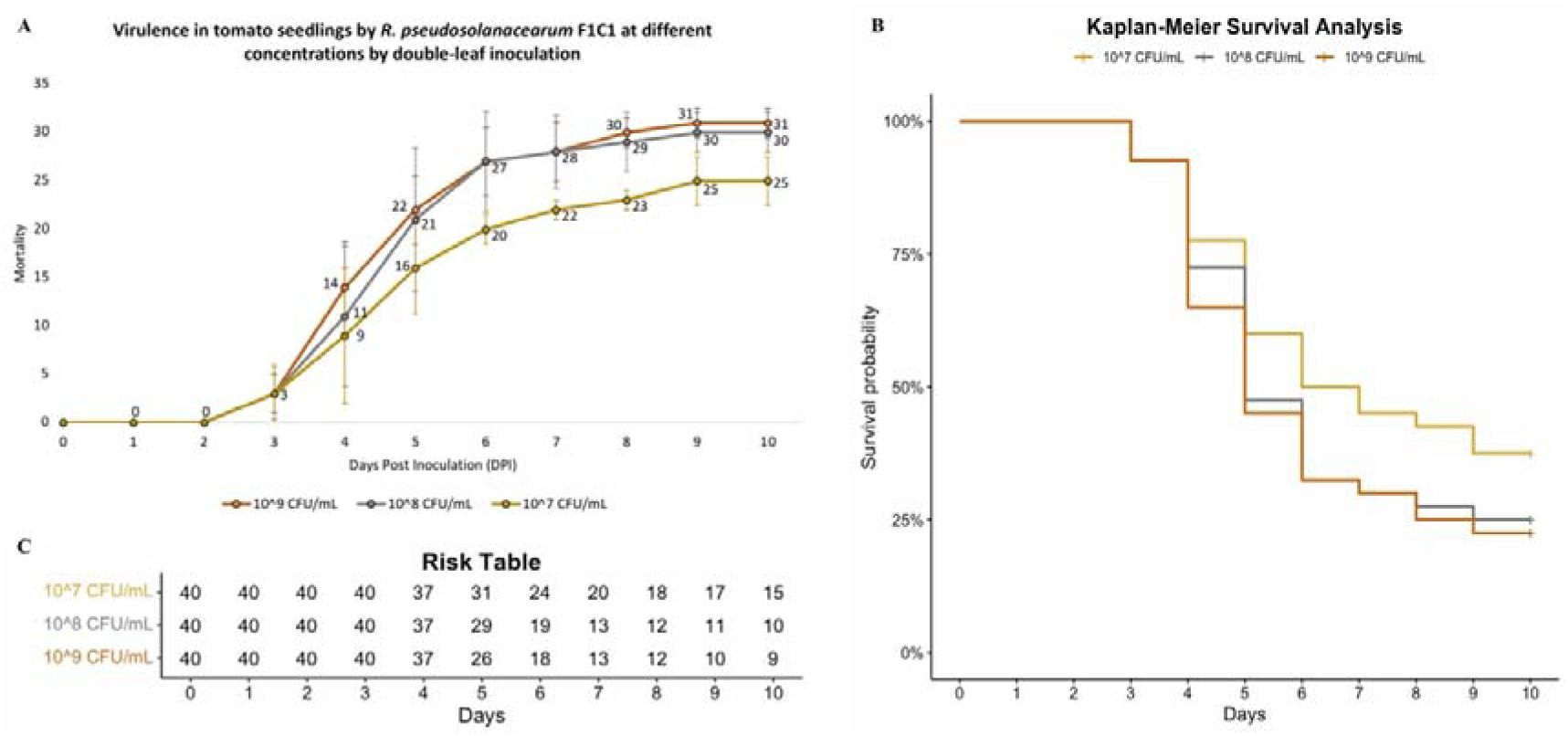
Double-leaf inoculation of higher concentrations (10^9^, 10^8^ and 10^7^ CFU/mL) of *R. pseudosolanacearum* F1C1 in tomato seedlings. **A.** Line-graph. **B.** Kaplan-Meier survival analysis. **C.** Risk Table.

A comparative study was done between the disease progression in tomato and eggplant at various concentrations of the pathogen such as 10^9^ CFU/mL, 10^8^ CFU/mL and 10^7^ CFU/mL using Log-rank analysis. At all the concentrations of the pathogen, the disease in eggplant was significantly higher than the disease in tomato (p-value < 0.05).

We studied further the pathogenicity of F1C1 by inoculating the eggplant and tomato seedlings at a lower concentration of the pathogen such as 10^6^ CFU/mL, 10^5^ CFU/mL, and 10^4^ CFU/mL (Fig. 4 & 5). In the case of eggplant, on 10 DPI the seedling mortality was 75 %, 57.5 %, and 10 %. Mortality of eggplant seedlings was initiated on 3 DPI at 10^6^ CFU/mL, while the same was initiated on 5 DPI at 10^5^ CFU/mL as well as 10^4^ CFU/mL. The disease progression between 10^6^ CFU/mL and 10^5^ CFU/mL was similar (p-value ≥ 0.05) while the disease progression at 10^4^ CFU/mL was significantly different (p-value < 0.05) from that of 10^6^ CFU/mL and 10^5^ CFU/mL. In the case of tomato seedlings, the seedling mortality was 47.5 %, 25 %, and 7.5 % at 10^6^ CFU/mL, 10^5^ CFU/mL, and 10^4^ CFU/mL respectively. In this case, the mortality was initiated on 5 DPI at 10^6^ CFU/mL as well as 10^5^ CFU/mL, and the same on 6 DPI at 10^4^ CFU/mL. The disease progression between 10^6^ CFU/mL and 10^5^ CFU/mL was similar (p-value ≥ 0.05) while the disease progression at 10^4^ CFU/mL was significantly different (p-value < 0.05) from that of 10^6^ CFU/mL and 10^5^ CFU/mL. The pathogenicity study in relation to pathogen load in the two hosts indicated that below a critical concentration of the pathogen, both the initiation, as well as incidence of the disease, are influenced by the pathogen load.

**Figure 4.**
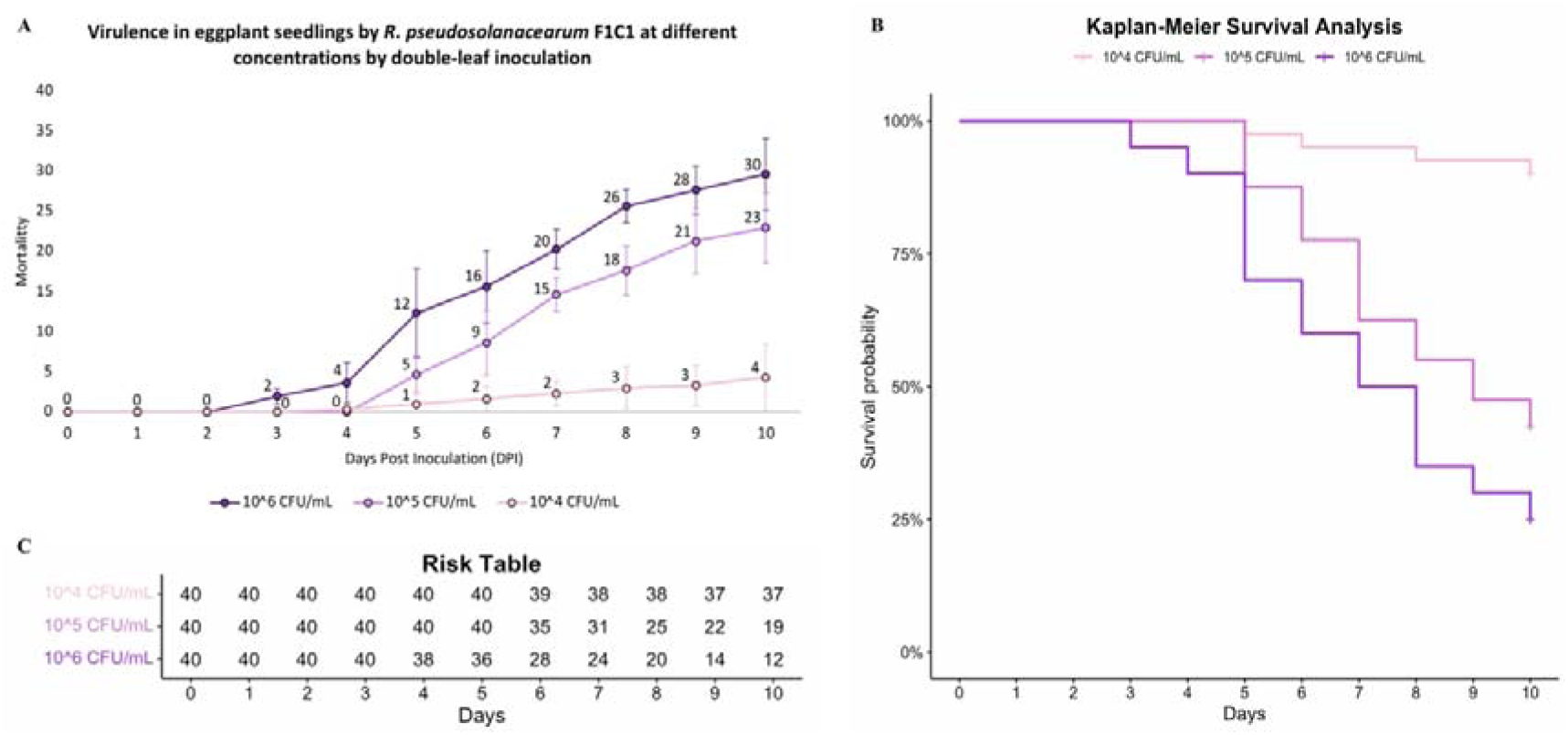
Double-leaf inoculation of lower concentrations (10^6^, 10^5^ and 10^4^ CFU/mL) of *R. pseudosolanacearum* F1C1 in eggplant seedlings. **A.** Line-graph. **B.** Kaplan-Meier survival analysis. **C.** Risk Table.

**Figure 5.**
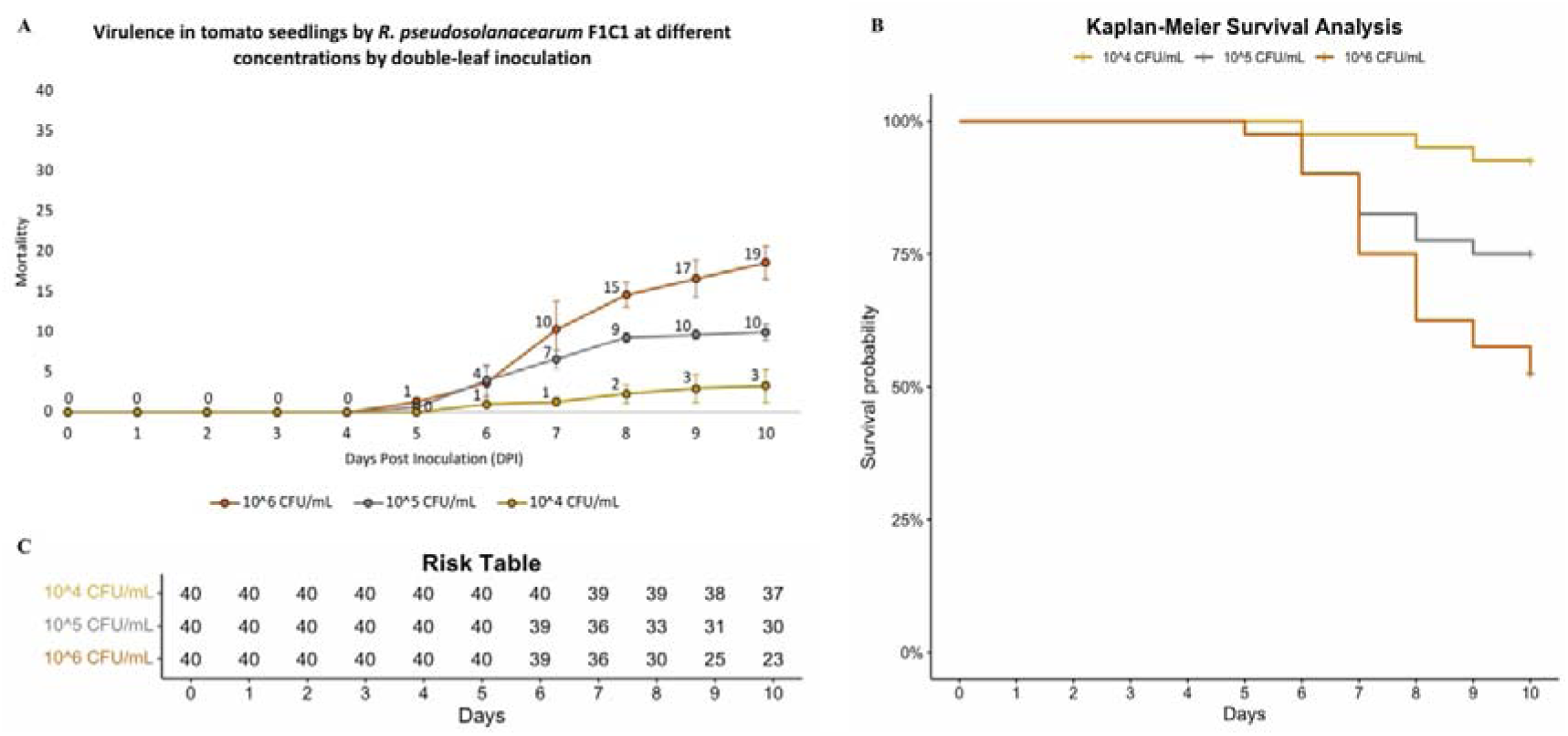
Double-leaf inoculation of lower concentrations (10^6^, 10^5^ and 10^4^ CFU/mL) of *R. pseudosolanacearum* F1C1 in tomato seedlings. **A.** Line-graph. **B.** Kaplan-Meier survival analysis. **C.** Risk Table.

The log-rank test was used to compare the impact of the pathogen load in the pathogenicity progression between tomato and eggplant. These results indicated that at pathogen concentrations of 10^6^ CFU/mL and 10^5^ CFU/mL, *R. pseudosolanacearum* exhibited significantly higher virulence in eggplant compared to tomato seedlings (p-value < 0.05). At a concentration of 10^4^ CFU/mL, there was no significant difference in virulence between the two plant types (p-value = 0.69) as expected because the incidences of the disease were low. Tomato and eggplant are phylogenetically close. Therefore, it was interesting to observe the distinct pathogenicity difference between the two hosts with regard to escapees observation, disease progression as well as pathogenicity difference regarding pathogen load by the leaf clip inoculation.

### (b) *R. pseudosolanacearum* F1C1 pathogenicity difference in tomato and eggplant seedlings using single and double-leaf inoculations

*R. pseudosolanacearum* causes a systemic infection under natural conditions. The pathogen enters the host plant from the soil through the root and then colonizes the shoot, ultimately leading to wilting. However, in the leaf clip inoculation study, the disease occurrence begins at the site of inoculation in the seedlings and then progresses downward. Both tomato and eggplant are dicots for which the seeds germinate to produce two cotyledon leaf seedlings. Therefore, it provided us with a novel avenue to do a comparative study of pathogenicity between the two hosts by single-leaf as well as double-leaf inoculations.

Eggplant seedlings were inoculated with the pathogen (10^9^ CFU/mL) by single-leaf as well as double-leaf inoculation and the impact of the two inoculation modes on pathogenicity was compared. Pathogenicity was observed on 3 DPI in both single- and double-leaf inoculations. However, on 4 and 5 DPI, a distinct difference in pathogenicity was evident between single- and double-leaf inoculations. To elaborate, with single-leaf inoculation the mortality rates on the 4 and 5 DPI were 7.5 % and 45 %, respectively, while with double-leaf inoculation, the mortality rates were notably higher at 60 % on the 4 DPI and 100 % on the 5 DPI. It is only by 8 DPI that 100 % mortality was also observed in the case of single-leaf inoculation. This experiment in eggplant proved a significant difference in pathogenicity between single- and double-leaf inoculations (p-value < 0.0001) (Fig. 6). The two modes of inoculation were further performed in eggplant seedlings at lower pathogen concentrations such as 10^8^ CFU/mL and 10^7^ CFU/mL (Fig. 7 & 8). As observed above, a significant difference (p-value < 0.05) in pathogenicity between single- and double-leaf inoculations was observed at these two concentrations. It is interesting to observe that double-leaf inoculation at 10^7^ CFU/mL exhibits similar pathogenicity to single-leaf inoculation at 10^8^ CFU/mL. Observations in this study supported the notion that within a host, inoculation at different sites results in higher pathogenicity than inoculation at a single site.

**Figure 6.**
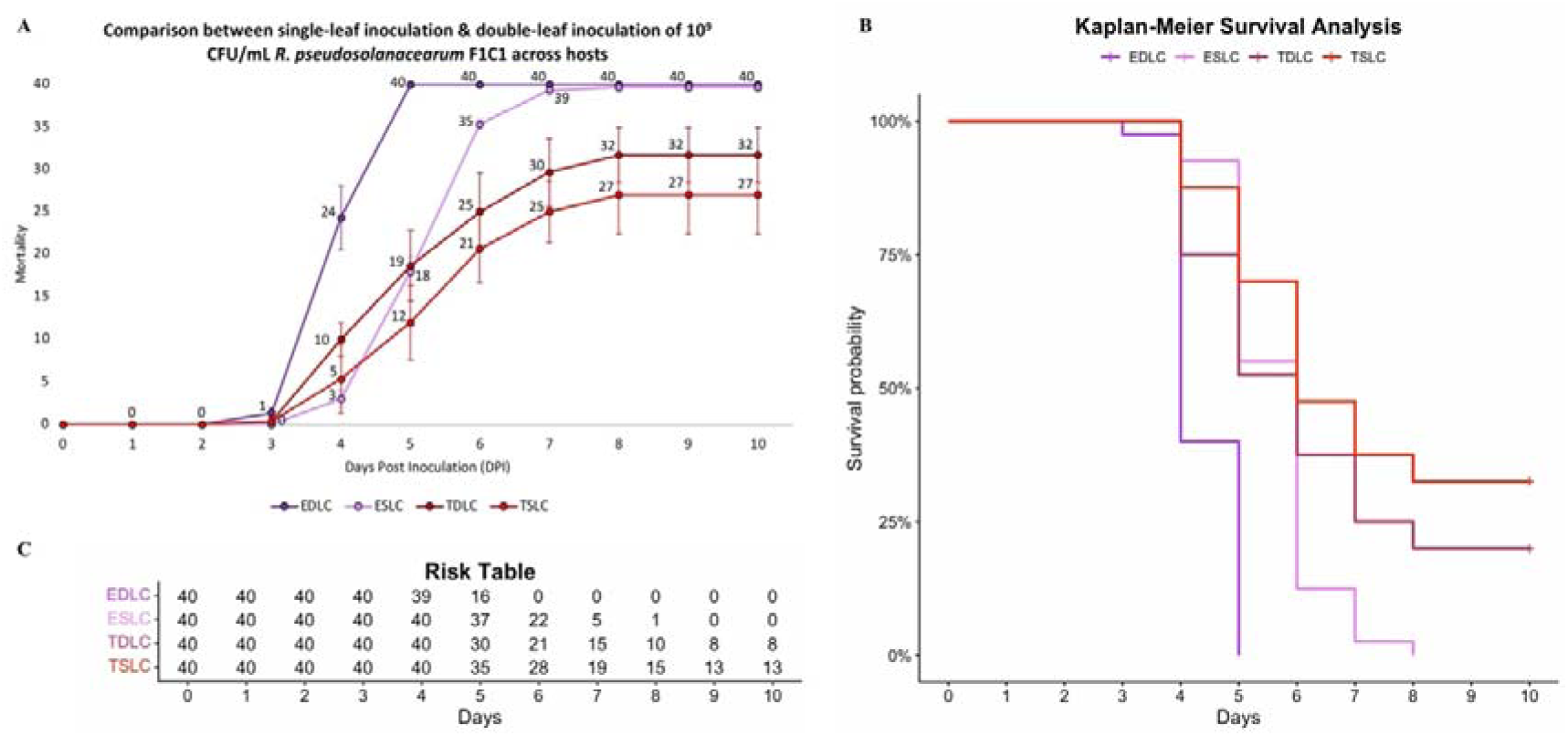
Single-leaf and double-leaf inoculation of 10^9^ CFU/mL *R. pseudosolanacearum* F1C1 in tomato and eggplant seedlings. **A.** Line-graph. **B.** Kaplan-Meier survival analysis. **C.** Risk Table. EDLC – Eggplant double-leaf clip; ESLC – Eggplant single-leaf clip; TDLC – Tomato double-leaf clip; TSLC – Tomato single-leaf clip.

**Figure 7.**
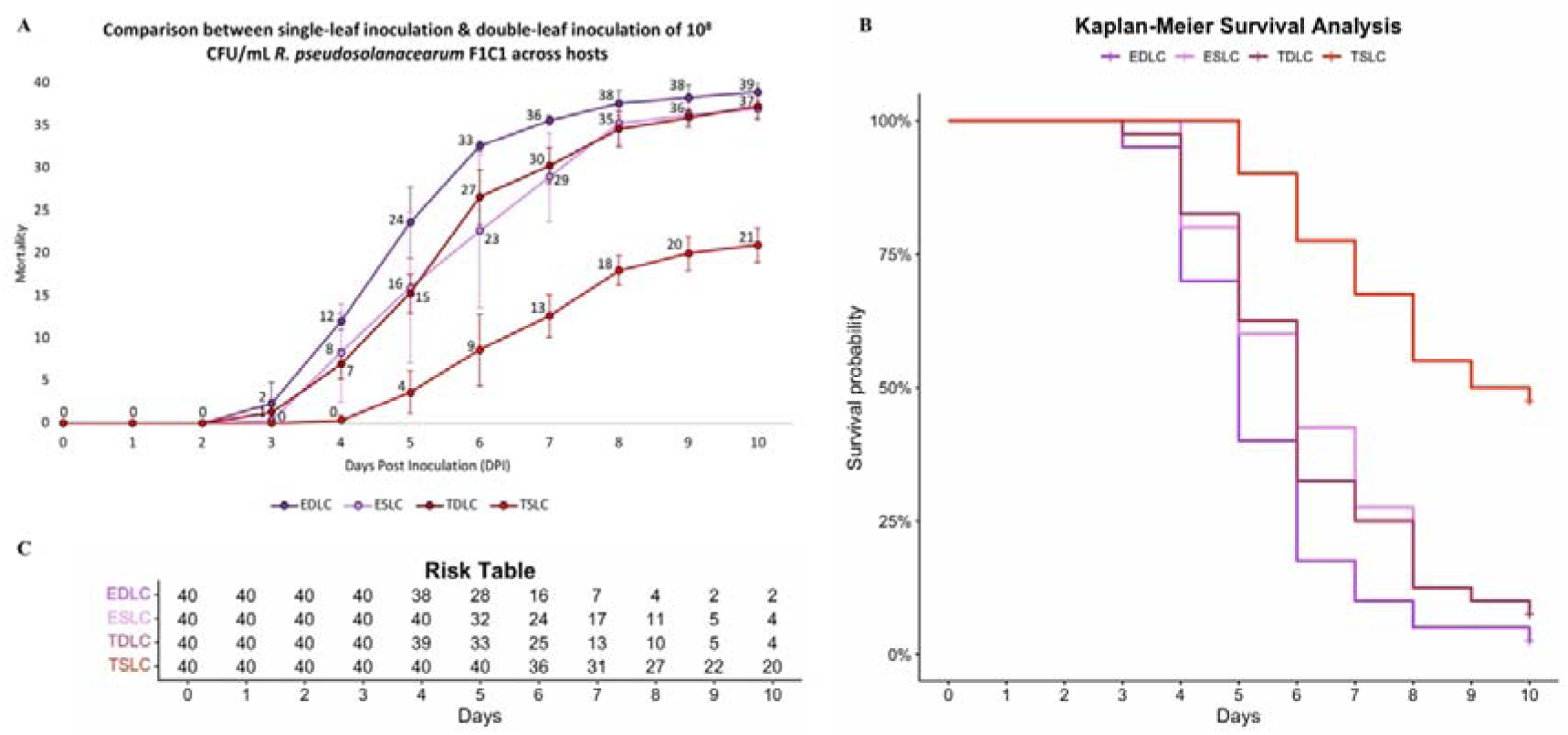
Single-leaf and double-leaf inoculation of 10^8^ CFU/mL *R. pseudosolanacearum* F1C1 in tomato and eggplant seedlings. **A.** Line-graph. **B.** Kaplan-Meier survival analysis. **C.** Risk Table. EDLC – Eggplant double-leaf clip; ESLC – Eggplant single-leaf clip; TDLC – Tomato double-leaf clip; TSLC – Tomato single-leaf clip.

**Figure 8.**
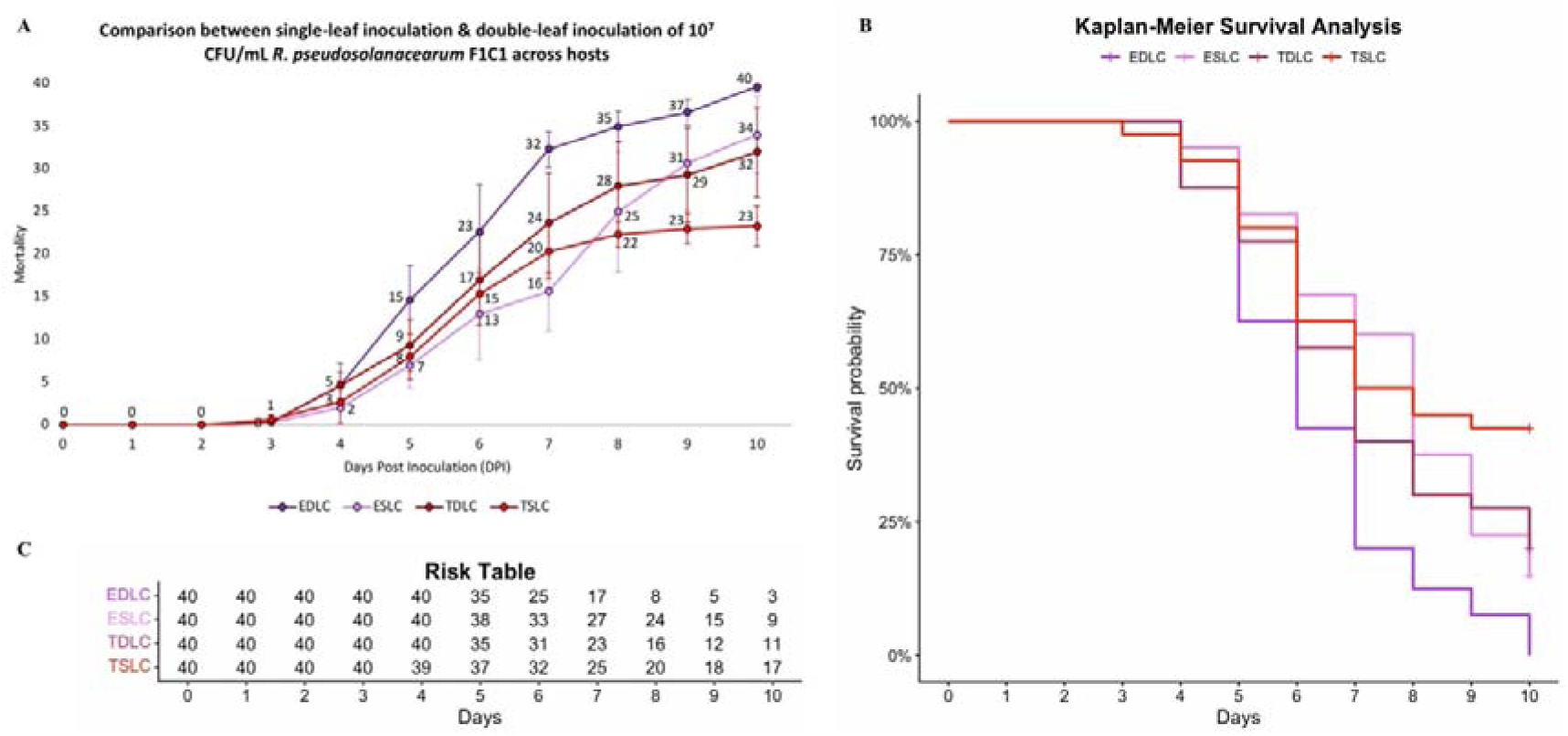
Single-leaf and double-leaf inoculation of 10^7^ CFU/mL *R. pseudosolanacearum* F1C1 in tomato and eggplant seedlings. **A.** Line-graph. **B.** Kaplan-Meier survival analysis. **C.** Risk Table. EDLC – Eggplant double-leaf clip; ESLC – Eggplant single-leaf clip; TDLC – Tomato double-leaf clip; TSLC – Tomato single-leaf clip.

Tomato seedlings were inoculated with the pathogen (10^9^ CFU/mL) by single-leaf as well as double-leaf inoculation and the impact of the two inoculation modes on pathogenicity was compared. The mortality of seedlings at 10 DPI were 80 % and 67.5 %, by double- and single-leaf inoculations, respectively (Fig. 6), and the pathogenicity difference was not significant (p-value > 0.05). The pathogenicity study in tomato seedlings by single- and double-leaf inoculations was further performed at lower pathogen concentrations, such as 10^8^ CFU/mL and 10^7^ CFU/mL (Fig. 7 & 8). It is indeed quite surprising to note that the single- or double-leaf inoculations in tomato seedlings did not result in any consistent difference in terms of pathogenicity, while the difference was distinct in the case of eggplant seedlings. This further demonstrated the difference between the two hosts and the advantage of studying pathogenicity in two different hosts.

### **(c)** Low susceptibility of eggplant seedlings than tomato seedlings to *R. pseudosolanacearum* infection by roots inoculation

*R. pseudosolanacearum* F1C1 pathogenicity in eggplant, as well as tomato seedlings by root inoculation, has already been demonstrated. Here we compared the two hosts regarding *R. pseudosolanacearum* F1C1 pathogenicity by root inoculation. Mortality in tomato seedlings was observed on 10 DPI as 87.5 % in 10^9^ CFU/mL, 75 % in 10^8^ CFU/mL and 57.5 % in 10^7^ CFU/mL (Fig. 9). Comparative statistical analysis through log-rank tests revealed no consistent difference in the intensity of virulence across the three different pathogen concentrations of 10^9^ CFU/mL, 10^8^ CFU/mL and 10^7^ CFU/mL. *R. pseudosolanacearum* F1C1 pathogenicity in eggplant seedlings was however found to be very low: only two seedlings (5 %) died out of 40 seedlings at 10^9^ CFU/mL (Fig. 10). This was consistently observed in the laboratory for which we did not study pathogenicity in eggplant seedlings by root inoculations at lower concentrations of the pathogen. This low pathogenicity in eggplant seedlings by root inoculation was further studied by the soil drenching method too where only five seedlings (12.5 %) died out of 40 seedlings at 10^9^ CFU/mL (Fig. 10, SFig. 2). In this study, it was observed that *R. pseudosolanacearum* pathogenicity is different between the two hosts such as tomato and eggplant by root inoculation.

**Figure 9.**
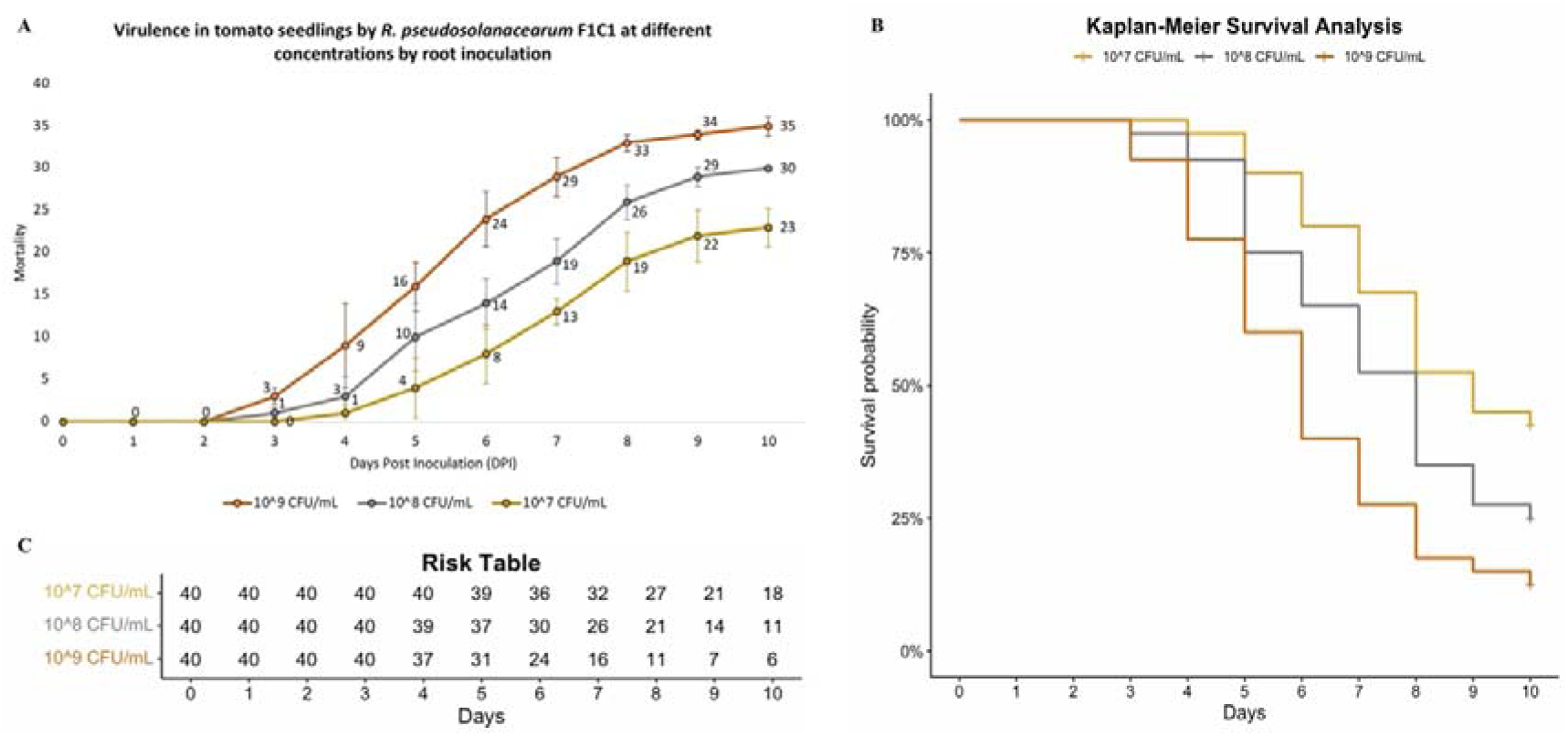
Root inoculation of different concentrations (10^9^, 10^8^ and 10^7^ CFU/mL) of *R. pseudosolanacearum* F1C1 in tomato seedlings. **A.** Line-graph. **B.** Kaplan-Meier survival analysis. **C.** Risk Table.

**Figure 10.**
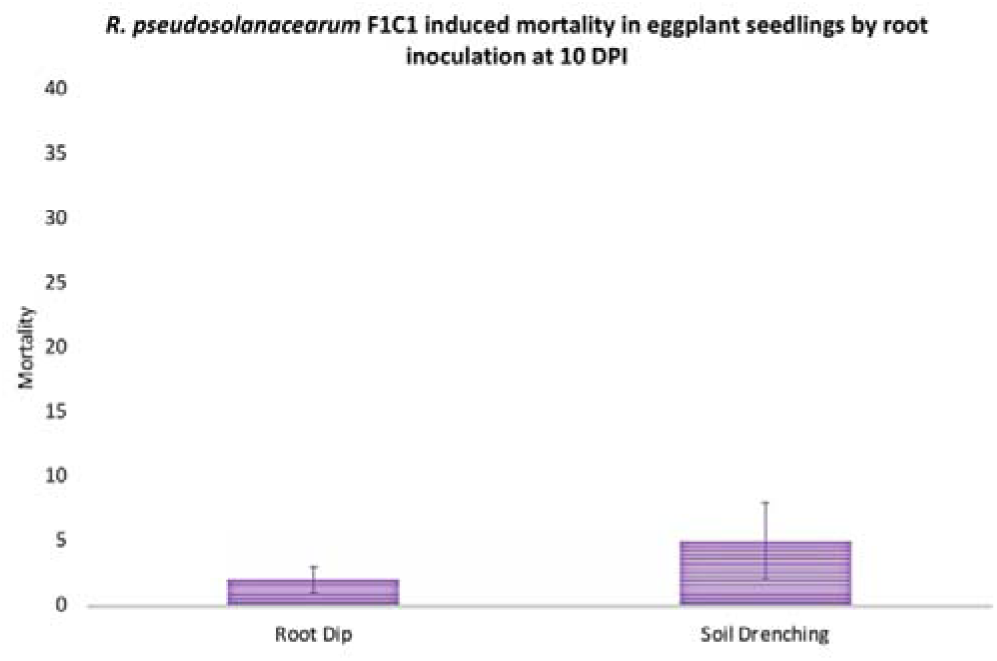
Mortality in eggplant seedlings at 10 DPI caused by root inoculation of *R. pseudosolanacearum* via root dip and soil drenching method.

## 4. Discussion

Albeit being anchored, plants exhibit directional movement in the form of positive phototropism and negative geotropism in their shoot region, but negative phototropism and positive geotropism in their root region. The tissues of leaves and roots are distinctive with regard to their physiology and metabolism, which allows the leaf tissues to perform photosynthesis in the presence of chlorophyll and the root tissues to absorb water and access nutrients. This plainly underscores the distinction between the two tissues. The present study demonstrates that these distinctions between the two tissues also extend to the plant’s susceptibility to pathogens. To empirically investigate this, *R. pseudosolanacearum*, a phytopathogen known to cause systemic wilting, was chosen as a model organism for the study rather than a phytopathogen causing localized disease.

*R. pseudosolanacearum* is a broad host range systemic phytopathogenic bacterium. However, under laboratory conditions, the tomato plant has primarily been used as the model host for assessing its pathogenicity. The use of multiple hosts is likely cumbersome considering the great demand for growing, maintaining and conducting virulence studies in grown-up plants. The use of seedlings in the virulence study of this pathogen has come up as a boon in this regard. Here, we have done a comparative pathogenicity study of *R. pseudosolanacearum* between two phylogenetically close solanaceous host seedlings of tomato and eggplant. Our observations suggest a distinct difference in virulence between the two host seedlings when inoculated by the cotyledon leaf, with eggplant seedlings demonstrating higher susceptibility than tomato seedlings. This higher susceptibility of eggplant than the tomato seedlings through leaf inoculations was further confirmed using single- and double-leaf inoculations. In the case of root inoculation in the seedlings, the tomato seedlings exhibited a greater susceptibility than the eggplant seedlings. This indicated that the susceptibility of the roots and the leaves in the same host to a systemic phytopathogen are independent of each other, which could only be revealed using this current comparative study involving two different hosts. Both in the case of tomato and eggplant, wilting frequency through the leaf inoculation is more than the root inoculation as cotyledon leaf inoculation provides direct access of the pathogen to the shoot region from where the disease symptom is observed. This is in concordance with the previously published work from the same laboratory [12].

The reason behind the higher susceptibility of the eggplant seedlings than the tomato seedlings through leaf inoculation and *vice versa* through root inoculation needs to be explored in future. The difference in the composition of the metabolites in the early stages of the cotyledon leaves between tomato and eggplant might be responsible for the differential susceptibility. Similarly, the metabolites present in the roots of the seedlings might be attributed towards their differential susceptibility. The differential pathogenicity of *R. pseudosolanacearum* F1C1 in the root and leaf tissues of the two hosts opens up prospects for the screening of host-specific virulence-deficient mutants through leaf as well as root inoculations.

The single- and double-leaf inoculations in seedlings have given us valuable insights into the plants’ defense response towards their pathogen. It sheds light on whether the host plant mounts an integrated systemic defence against a pathogen at the infection site or its defence response to pathogen infection primarily operates on a local level. In the case of double-leaf inoculations, wounds were created at two sites concurrently with pathogen introduction. In contrast, single-leaf inoculation involved creating wounds at two sites but pathogen introduction at only one site. Despite repeating the experiments with tomato seedlings multiple times with different researchers, there was no distinct difference regarding virulence between double-leaf inoculations and single-leaf inoculations. This may be attributed to the occurrence of escapees in tomato seedlings, which were common in our trials. Understanding why escapees occur more frequently in tomato seedlings could be a valuable area for future research, especially with a highly susceptible tomato variety. Our findings suggest that pathogen infection at one site does not significantly influence infection at the other site. If there were such an influence, we would expect the disease incidence in single-leaf inoculations to be notably lower than in double-leaf inoculations. Additionally, the instances of partial wilting also favor an explanation of the disease being a localized phenomenon rather than affecting the plant in its entirety. In the case of eggplant seedlings, double-leaf inoculations resulted in faster wilting than single-leaf inoculations, likely due to the higher susceptibility of eggplant cotyledon leaves to the pathogen. Therefore, host susceptibility is likely the primary factor responsible for the disease phenotype rather than the pathogen’s discrimination between the two hosts of tomato and eggplant.

## Ethics

Our work does not contain experiments involving animals and/or human participants.

## Data accessibility

The article has no additional data.

## Declaration of AI use

We have not used AI-assisted technologies in creating this article.

## Authors’ contributions

**S.B.:** Investigation, methodology, formal analysis, software, validation, visualisation, writing-original draft, writing-review and editing; **M.J.:** Investigation, methodology, visualisation, validation, writing-review and editing; **L.B./T.S.T.:** Investigation, methodology, validation,; **S.Begum/LD/SJG:** Investigation, validation, writing-review and editing; **T.P./K.K.:** Investigation, methodology, writing-review and editing; **M.M.:** Supervision, writing-review and editing; **S.K.R.:** Conceptualization, funding acquisition, supervision, visualisation, writing-original draft, writing-review and editing.

## Acknowledgements

**S.B.** is thankful for the JRF/SRF fellowship from the UGC-NFSC, GoI, New Delhi.
**M.J.** is thankful to Tezpur University for the institutional fellowship. **L.D.** is thankful to DST, GoI, New Delhi for the INSPIRE-JRF fellowship. **S.J.G.** and **S.Begum** are thankful for the JRF fellowship from the DBT, GoI New Delhi grant (BT/PR41637/NER/95/1753/2021) awarded to **S.K.R. S.K.R.** is thankful to his former collaborators Drs. Rahul Kumar, A. Barman, N. Singh, P. L. Sharma and N. Agarwala for the critical discussions on leaf inoculations.

