## Supplementary data for "Tissue-dependent nature of plant susceptibility: a comparative pathogenicity study of the systemic phytopathogen *Ralstonia pseudosolanacearum* in eggplant and tomato seedlings through root and leaf"

**Supplementary Figure 1.** Screenshot of a portion of the RStudio terminal showing the R script used for plotting the Kaplan-Meier survival curve and performing the Log-rank test.

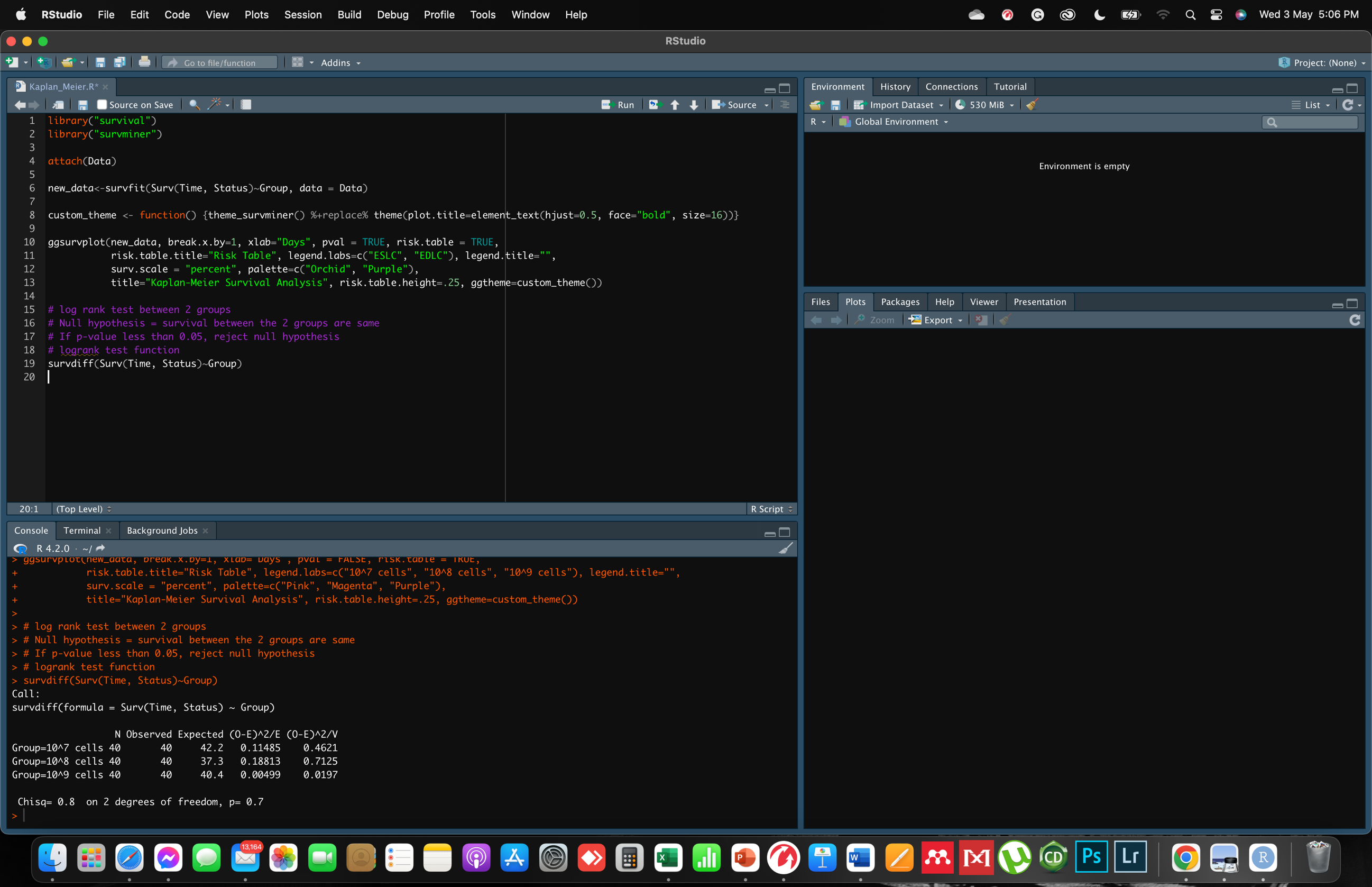

**Supplementary Figure 2.** Disease progression in eggplant seedlings inoculated with *R. pseudosolanacearum* F1C1 using the soil-drenching method.

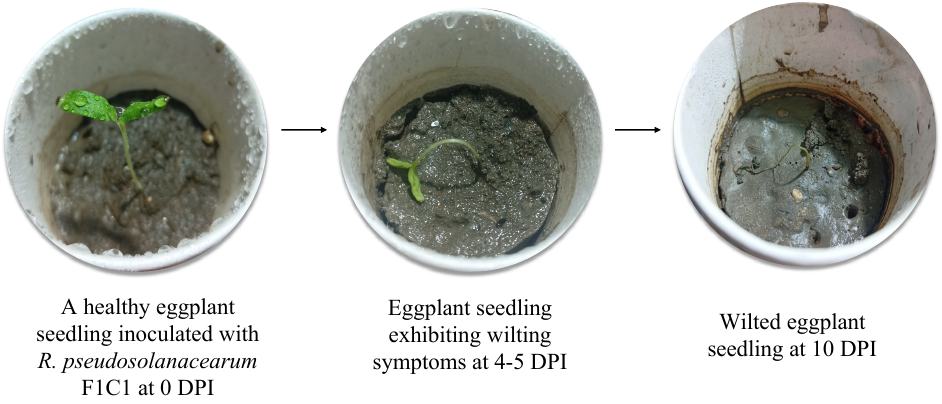
